# Optimal feature selection and software tool development for bacteriocin prediction

**DOI:** 10.1101/2022.09.29.510068

**Authors:** Suraiya Akhter, John Miller

## Abstract

Antibiotic resistance is a major public health concern around the globe. As a result, researchers always look for new compounds to develop new antibiotic drugs for combating antibiotic-resistant bacteria. Bacteriocin becomes a promising antimicrobial agent to fight against antibiotic resistance, due to its narrow killing spectrum. Sequence matching methods are widely used to identify bacteriocins by comparing them with the known bacteriocin sequences; however, these methods often fail to detect new bacteriocin sequences due to sequences’ high diversity. The ability to use a machine learning approach can help find new highly dissimilar bacteriocins for developing highly effective antibiotic drugs. The aim of this work is to identify optimal sets of features and develop a machine learning-based software tool for predicting bacteriocin protein sequences with high accuracy. We extracted potential features from known bacteriocin and non-bacteriocin sequences by considering the physicochemical and structural properties of the protein sequences. Then we reduced the feature set using statistical justifications and recursive feature elimination technique. Finally, we built support vector machine (SVM) and random forest (RF) models using the selected features and our models can achieve accuracy up to 95.54%. We compared the performance of our method with a popular sequence matching-based approach and a deep learning-based method. We also developed a software tool called Bacteriocin Prediction (BacPred) that implements the prediction model using the optimal set of features obtained from this study. The software package and its user manual are available at https://github.com/suraiya14/ML_bacteriocins/BacPred.

## Introduction

Bacteria become antibiotic resistant due to the excessive use of drugs in healthcare and agriculture. In the United States, around 3-million people get infected and approximately 35000 individuals die because of antibiotic-resistant organisms [1]. Therefore, the resistance nature of bacteria drives the need for inventing novel antimicrobial compounds to treat antibiotic-resistant patients. Researchers developed several approaches to extract natural products as antimicrobial compounds by mining the bacterial genomes [2]. Bacteriocin is one type of natural antimicrobial compound which is a bacterial ribosomal product. Due to narrow killing spectrum, bacteriocins became attractive choices in the discovery of novel drugs that can produce less resistance in bacteria [3–5]. Current whole genome sequencing technology provides many genes that encode bacteriocins and these sequences are publicly available for future research. Researcher introduced several methods to identify bacteriocins from bacterial genomes based on bacteriocin precursor genes or context genes. For example, BAGEL [6] and BACTIBASE [7] are two publicly available online tools that curate huge experimentally validated and annotated bacteriocins. Like the widely used protein searching tool BLASTP [8, 9], these methods also allow users to identify putative bacteriocin sequences based on the homogeneity of known bacteriocins. However, these similarity-based approaches often fail to detect sequences that have the high dissimilarity with known bacteriocin sequences; thereby, generating an undesired number of false negatives. To resolve this problem, some prediction tools, such as BOA (Bacteriocin Operon Associator) [10], were developed based on locating conserved context genes of the bacteriocin operon, but they still rely on homology-based genome searches.

Machine learning technique can be applied as a substitute for sequence similarity and context-based methods that can utilize potential peptide (protein) features of bacteriocin and non-bacteriocin to make strong prediction in identifying novel bacteriocin sequences. Recently some machine learning-based bacteriocin prediction techniques were proposed that utilized the presence or absence of *k*-mer (i.e., subsequences of length *k*) as potential features and represented peptide sequences using word embedding [11, 12]. There are also deep learning-based methods for bacteriocin prediction, for example RMSCNN [13] used a convolutional neural network [14, 15] for identifying marine microbial bacteriocins. However, these existing approaches did not consider the primary and secondary structure information of peptides that are crucial to find highly dissimilar bacteriocins. Also, those strategies did not apply any feature evaluation algorithm to eliminate the unnecessary features that may reduce the achievement of a machine learning classifier.

In this work we present a predictive pipeline for identifying bacteriocins by generating features from the physicochemical and structural characteristics of peptide sequences. We evaluated and selected subsets of the candidate features based on Pearson correlation coefficient, *t*-test, mean decrease Gini (MDG), and recursive feature elimination (RFE) analyses. The reduced feature sets called optimal feature sets are then used to predict bacteriocins using support vector machine (SVM) [16, 17] and random forest (RF) [18] machine learning models. BLASTP, a sequence matching tool and RMSCNN, a deep learning model were used to compare the performance of our best machine learning model. One of the main objectives was to develop a software package called Bacteriocin Prediction (BacPred) with a simple and intuitive graphical user interface (GUI) that can generate all required features to get prediction results for testing protein sequences. The software provides options to users to test multiple sequences and add new training bacteriocin or non-bacteriocin sequences to the machine learning model for improving the prediction capability.

## Materials and methods

The overall workflow of our methods is depicted in Fig 1. The steps in our methods include gathering datasets of bacteriocin and non-bacteriocin protein sequences, generating potential features, performing feature evaluation and recursive feature elimination analyses to remove irrelevant and weakest features, and finally building machine learning models using the selecting features to compare the prediction performance with the sequence matching and deep learning-based approaches.

**Fig 1.**
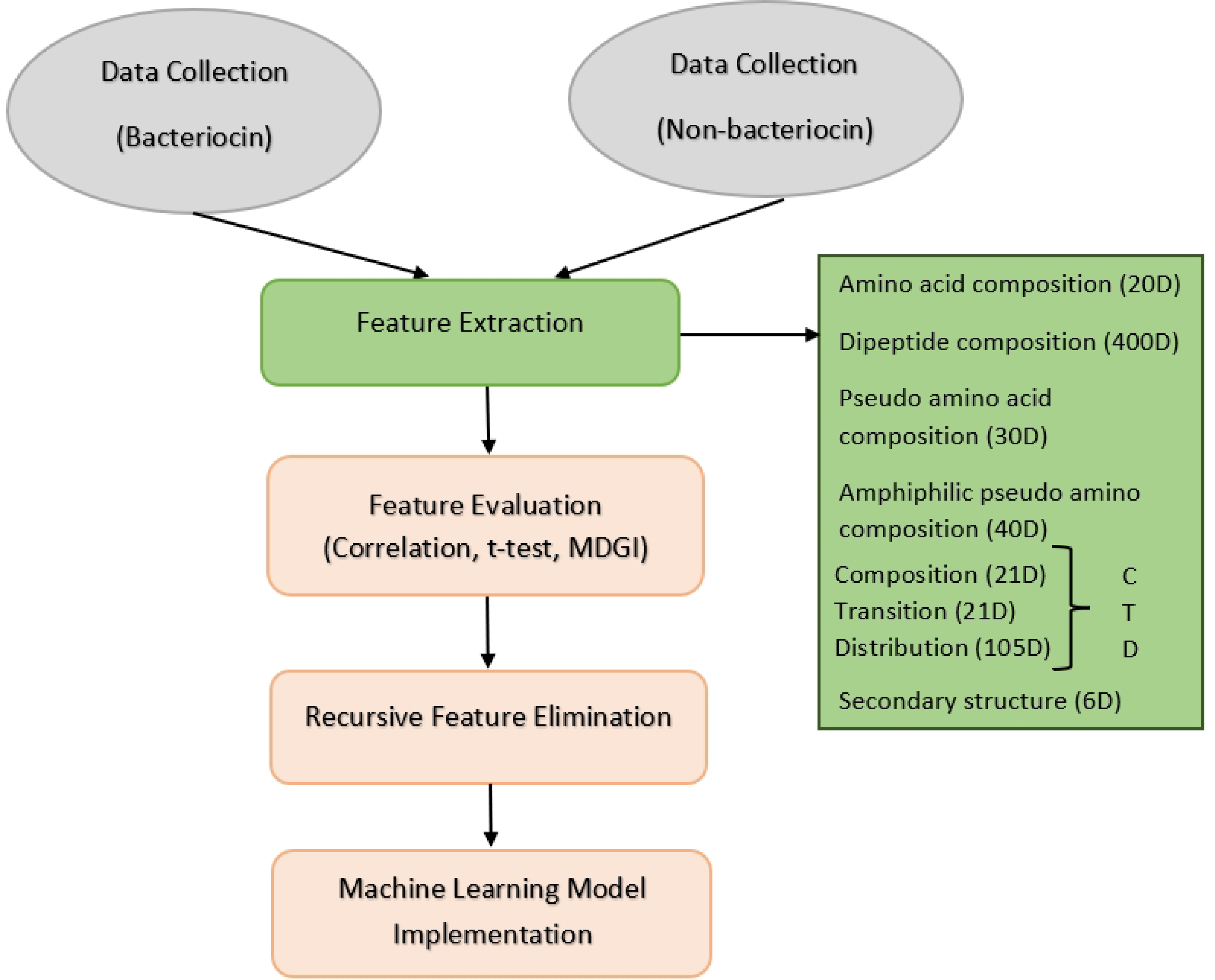
Illustrating the steps involved in the prediction of bacteriocin protein sequences.

### Data collection

We retrieved bacteriocin sequences (positive sequences) from two publicly available databases BAGEL [6] and BACTIBASE [7]. Non-bacteriocin sequences (negative sequences) were collected from the data used in RMSCNN [13]. Initially, we gathered a total of 483 positive and 500 negative sequences. As many of these accumulated sequences are duplicate or of high similarity, we utilized CD-HIT tool [19] to obtain the unique positive and negative sequences by removing the sequences having ≥90% similarity. Finally, we obtained 283 and 497 unique positive and negative sequences, respectively. To deal with the imbalanced dataset problem, we reduced the negative sequences from 497 to 283 by random sampling to make the number of positive and negative examples equal. We considered 80% and 20% of the total sequences as training and testing datasets, respectively. Positive and negative training sequences, in FASTA format, are listed in **S1 File**. Positive and negative testing sequences are presented in **S2 File**.

### Feature extraction

After collecting the positive and negative protein sequences, we generated potential candidate features from the sequences. Since there are 20 natural amino acids, we generated a 20D (‘D’ indicates dimension) amino acid composition (AAC) feature vector for every protein sequence where each value in the vector gives the fraction of a specific amino acid type. We extracted 400D dipeptide composition (DC) feature vectors for the sequences where each value indicates the fraction of dipeptides in a protein sequence [20]. Pseudo amino acid composition (PseAAC) and amphiphilic pseudo amino acid composition (APseAAC) feature vectors of 30D and 40D, respectively, were created for each sequence as proposed by Chou [21, 22].

We used the composition/transition/distribution (CTD) model [23] to generated 147D feature vectors for various physicochemical amino acid properties. Amino acids are divided into three classes in the CTD model. For each sequence, we obtained 3D, 3D and 15D feature vectors for each physicochemical property as measurements of the composition, transition, and distribution of the classes, respectively. Finally, we generated 6D feature vectors from the secondary structure (SS) of each sequence. The SS features includes position index, spatially consecutive states, and segment information of the 3 structure states: alpha helices, beta sheets and gamma coils. In total, we obtained a total of 643 features as listed in Table 1.

**Table 1.** List of features.

| Feature | Dimension |
| --- | --- |
| AAC | 20 |
| DC | 400 |
| PseAAC | 30 |
| APseAAC | 40 |
| CTD | 147 |
| SS | 6 |

### Feature evaluation

Unnecessary features may worsen the prediction performance of a machine learning model. We performed statistical analyses on the training data to identify the optimal or best feature sets to build our machine learning models. At first, we estimated Pearson correlation coefficient *ρ*_*x,y*_ given by Eq (1), to measure the correlation values among features.

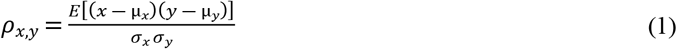

Here, *x* and *y* are two features, *E* indicates the expectation, *σ*_*x*_ and *σ*_*y*_ indicate the standard deviation, and μ_*x*_ and μ_*y*_ are mean values of *x* and *y*, respectively. High absolute the value of *ρ*_*x,y*_ indicates strong correlation with other features. If a feature is highly correlated with another feature, we can consider one of these two features and ignore the other one. We removed one of the two features if they have correlation value was ≥0.9, which resulted in the number of features decreasing from 643 to 590.

Then we considered two additional statistical approaches to feature reduction. First, a standard *t*-test was applied to each of the 590 features to see if a statistical significant difference existed between the values of the feature in the positive and negative bacteriocin sequences of our dataset. We estimated the *p*-values for all 590 features to check if it was possible to discard the null hypothesis of no statistically significant difference. A low *p*-value for a feature indicates high importance of the feature for predicting bacteriocin sequences, and in that situation, we can discard the null hypothesis. We considered a threshold *p*-value of 0.05 and eliminated all features having *p* > 0.05. After filtering the features based on the *t*-test results, our feature vector was reduced from 590D to 140D, and we called the resulting data the *t-test-reduced feature set*. The *p*-values of the selected features are shown in Fig 2 on linear and logarithmic scales. We noticed that the composition and distribution features of the CTD model were the top selected features in the *t*-test-reduced feature set.

**Fig 2.**
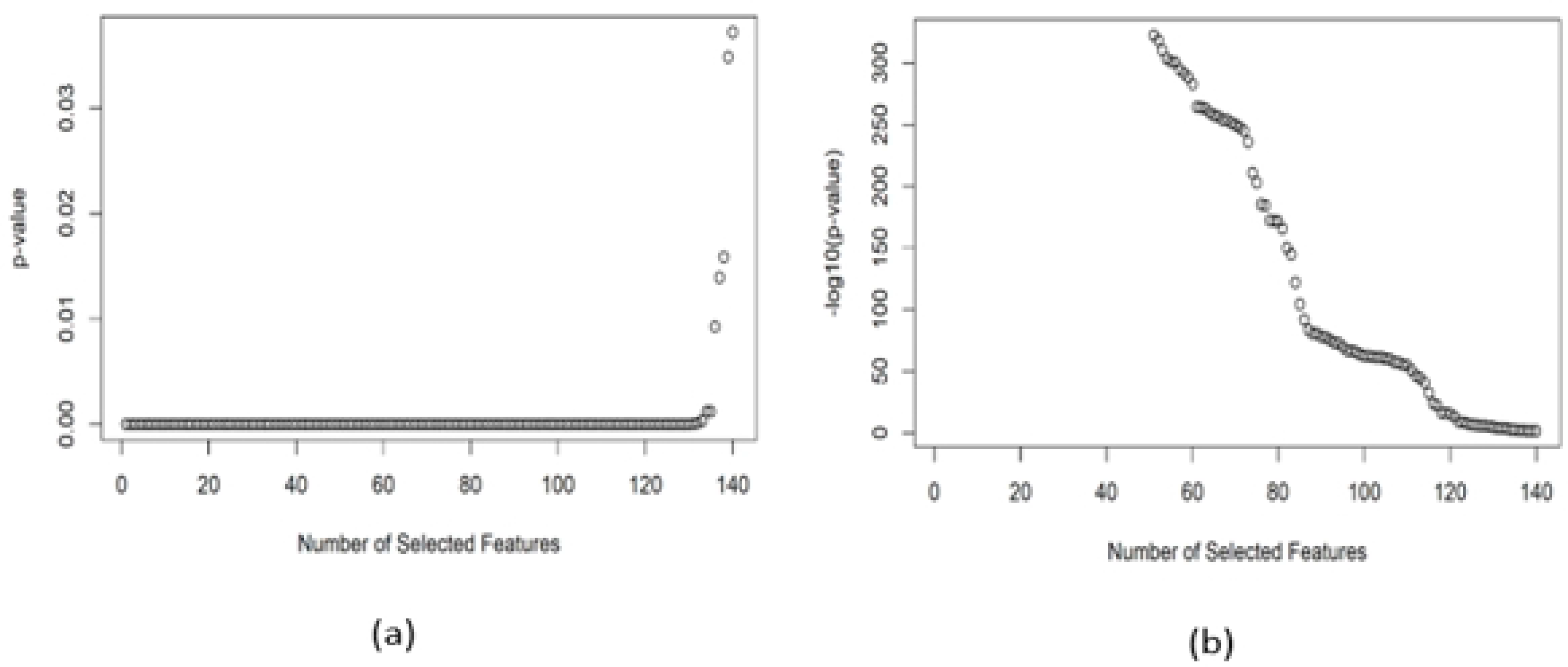
Trends of the *p*–values of the reduced feature set: (a) *p*–value vs. selected features and (b) −*log*_*10*_(p-value) vs. selected features.

Lastly, we built the random forest (RF) model with the 590 features (obtained from the Pearson correlation coefficient analysis) to estimate the mean decrease Gini (MDG). In the RF model, MDG corresponds to the feature importance that indicates each feature’s contribution to the homogeneity of the nodes and leaves [24, 25]. Eq (2), where *P*_*i*_ is the probability of being in class *i* (positive or negative), was used to calculate the Gini index. A node is purer when its Gini index is closer to 0.

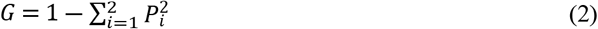

Gini index of 0 and 1 corresponds to complete homogeneity and heterogeneity of the data, respectively. MDG is computed from the mean of all the drop of Gini indices across the whole of the trees built in the RF model. Greater MDG value indicates a more important feature, and with consideration of MDG values for the features, we reduced the dimension of the feature set to 44D and named it the *MDG-reduced feature set*. Features of the CTD model, PseAAC, and SS were identified as top selected features in the MDG-reduced feature set.

### Recursive feature elimination

We further filtered features from the *t*-test-reduced and MDG-reduced feature sets using the recursive feature elimination (RFE) technique where a machine learning model is fitted, and features were ranked based on the evaluation of the training performance of the model. We considered two machine learning models RF and SVM in the RFE analyses and we obtained 42 (RF with MDG-reduced feature sets), 57 (RF with *t*-test-reduced feature sets), 44 (SVM with MDG-reduced feature sets) and 131 (SVM with *t*-test-reduced feature sets) features. We applied 5 times repeated 10-fold cross-validation to assess the capability of the SVM and RF in the training phase in the RFE analyses.

### BacPred software tool

Fig 3 shows the GUI of our BacPred software tool. All the required features in the BacPred tool were generated using R and the GUI was designed using Python3. In this tool, users can upload and save input file that should contain all protein sequences in FASTA format. If a user chooses the option of predicting bacteriocin, the BacPred software tool will consider all protein sequences in the input file as testing sequences and generate all required features with their feature values for the testing protein sequences automatically, classify them as bacteriocin or non-bacteriocin sequences and save the results in an output file. Users can add new bacteriocin or non-bacteriocin protein sequences to the training dataset and return to the original training dataset supplied with this tool, if desired. The tool has a textbox in the GUI where users can see the prediction results. The software package and the manual to use this software can be found at https://github.com/suraiya14/ML_bacteriocins/BacPred.

**Fig 3.**
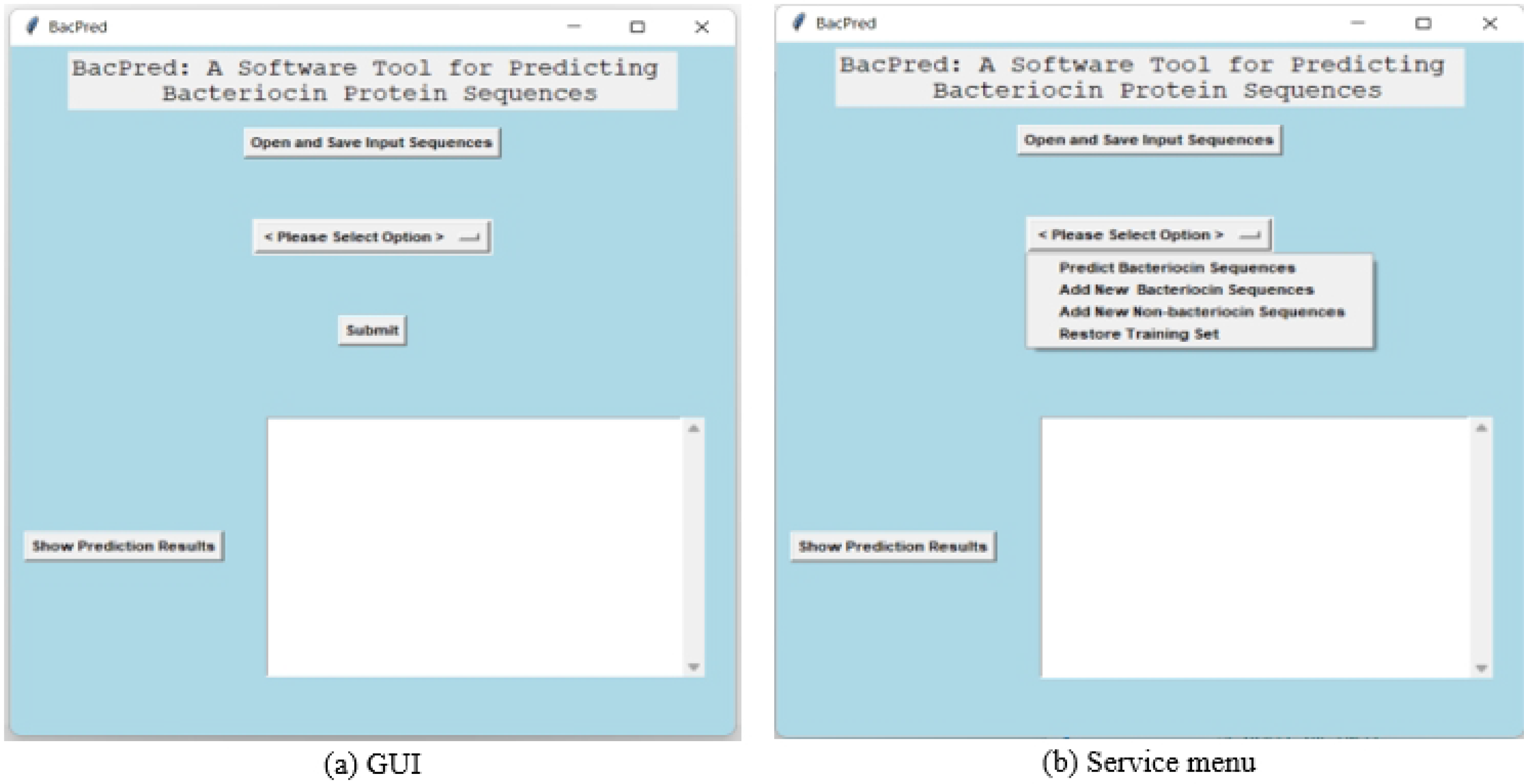
Graphical user interface (GUI) and various service menus of the BacPred software tool.

## Code and data availability

All experimental data and scripts of this work are available at https://github.com/suraiya14/ML_bacteriocins.

## Results and discussion

We measured the performance of SVM and RF machine learning models with the selected feature sets. We also performed a comparative analysis with sequence-matching and deep-learning approaches.

### Feature rankings

We mentioned earlier that SVM and RF machine learning models were used in the RFE approach to measure the training performance in terms of area under the ROC curve (AUC) by recursively considering subsets of the *t*-test-reduced and MDG-reduced feature sets independently. Figs 4(a) and 4(b) show the AUC values for the subset of the features in the RFE approach where RFE-MDG-RF and RFE-MDG-SVM depict the RFE analyses with the MDG-reduced feature sets for RF and SVM machine learning models, respectively. Similarly, Figs 4(c) and 4(d) are RFE analyses with the *t*-test-reduced feature sets for RF and SVM machine learning models, respectively. We noticed gradual decreasing of AUC values with the elimination of the features from the machine learning models. Table 2 lists the maximum AUC values obtained from the machine learning models in the RFE analyses. We obtained highest AUC value in the RF model for the MDG-reduced feature set.

**Table 2.**
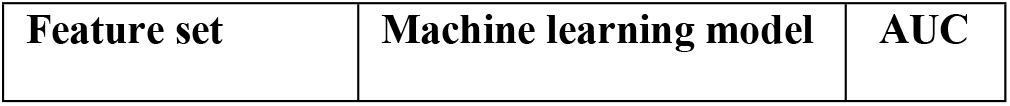
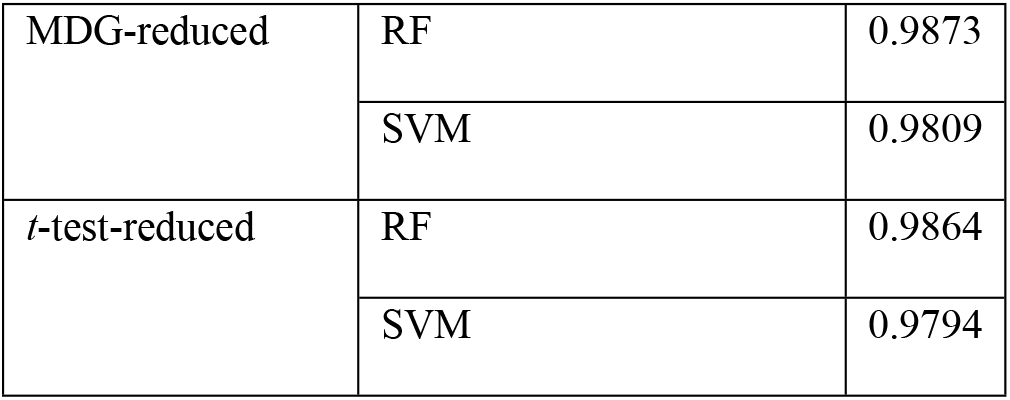
Highest AUC values obtained from RF and SVM for different feature sets.

| Feature set | Machine learning model | AUC |
| --- | --- | --- |
| MDG-reduced | RF | 0.9873 |
|  | SVM | 0.9809 |
| *t*-test-reduced | RF | 0.9864 |
|  | SVM | 0.9794 |

**Fig 4.**
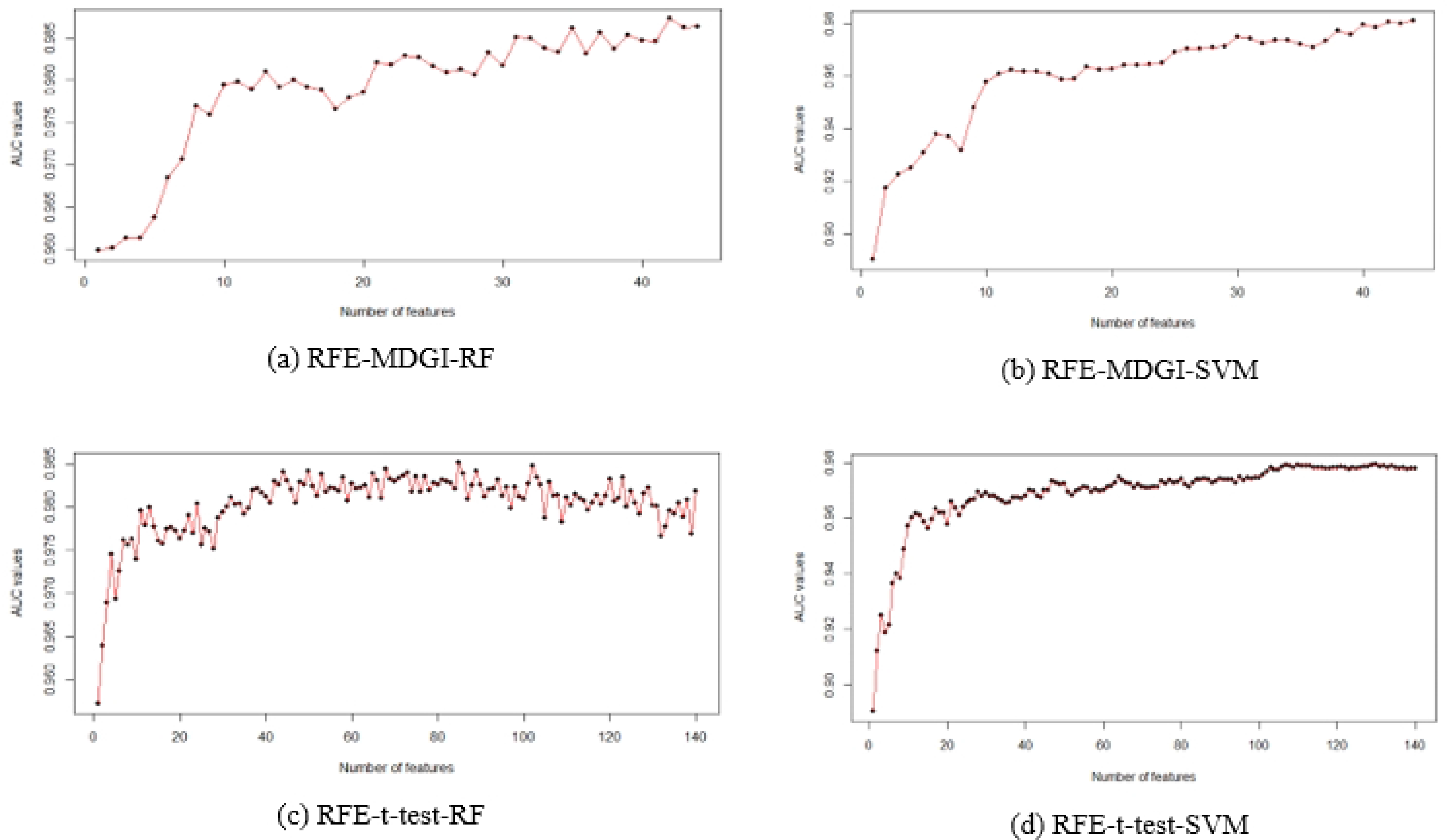
Performance of the RF and SVM machine learning models for the training data in the RFE approach.

The top-5 features obtained from the RFE analyses are listed in Table 3. Features of the CTD model and PseAAC features are among the top ranked features for all models. More specifically, distribution (first residue) for secondary structure (group 1), distribution (first residue) for hydrophobicity (group 3) and distribution (first residue) for normalized van der Waals Volume (group 3) of the CTD model were found common in the top-5 features of all RFE analyses.

**Table 3.** Top ranked features found from RF and SVM models in the RFE analyses.

| Feature rank | Feature for RFE-MDG-RF | Feature for RFE-MDG-SVM | Feature for RFE-*t*-test-RF | Feature for RFE-*t*-test-SVM |
| --- | --- | --- | --- | --- |
| 1 | Distribution (first residue) for hydrophobicity (group 3) | PseAAC for the amino acid Leucine (L) | Distribution (first residue) for charge (group 2) | PseAAC for the amino acid Leucine (L) |
| 2 | Distribution (first residue) for secondary structure (group 1) | PseAAC for the amino acid Arginine (R) | Distribution (first residue) for hydrophobicity (group 3) | PseAAC for the amino acid Arginine (R) |
| 3 | Distribution (first residue) for charge (group 2) | Distribution (first residue) for hydrophobicity (group 3) | Distribution (first residue) for solvent accessibility (group 3) | Distribution (first residue) for hydrophobicity (group 3) |
| 4 | Distribution (first residue) for solvent accessibility (group 3) | Distribution (first residue) for secondary structure (group 1) | Distribution (first residue) for secondary structure (group 1) | Distribution (first residue) for secondary structure (group 1) |
| 5 | Distribution (first residue)<br>for normalized van der<br>Waals Volume (group 3) | Distribution (first residue) for<br>normalized van der Waals<br>Volume (group 3) | Distribution (first residue)<br>for normalized van der<br>Waals Volume (group 3) | Distribution (first residue)<br>for normalized van der<br>Waals Volume (group 3) |

### Prediction performance

For our reduced feature sets, we trained SVM and RF models with different feature subsets obtained after RFE analyses. We tuned the SVM model with radial basis function (RBF) and set of cost values *C* = {4, 8, 16, 32, 64, 128} to find the best parameters. Similarly, we tuned the RF model with setting *ntree* = {400, 500} and *mtree* = {5, 6}. The RBF-kernel SVM with cost values of 4, 4, 4 and 8, and RF with *ntree* values of 500, 400, 500 and 400 and *mtree* values of 6, 5, 6 and 6 were found as best parameters for RFE-MDG-RF, RFE-MDG-SVM, RFE-*t*-test-RF and RFE-*t*-test-SVM feature sets, respectively.

To find the best optimal feature set, we measured test performance of our tuned models, SVM and RF, for the reduced feature sets. We evaluated the prediction performance using Eqs (3) and (4), where TP, TN, FP, and FN correspond to true positives (correctly classified as positives values), true negatives (correctly classified as negative values), false positives (incorrectly classified as positive values), and false negatives (incorrectly classified as negative values), respectively. The Matthews correlation coefficient (MCC) [26, 27] estimation is considered to measure the effectiveness of our classifiers and the MCC value ranges from −1 to +1. The larger the MCC value, the better prediction is.

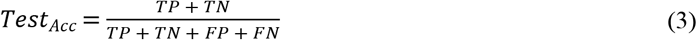

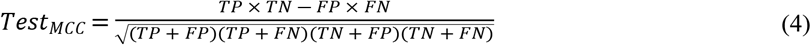

The prediction results of the models with corresponding best parameters are shown as confusion matrices in **S1-S8 Tables** where ‘1’ and ‘−1’ indicate positive (bacteriocin) and negative (non-bacteriocin) sequences, respectively. The diagonal entries show the correctly classified test sequences. The testing MCC and accuracy values (indicated as *Test*_*MCC*_ and *Test*_*ACC*_, respectively) of the RF and SVM models for different feature subset after RFE analyses are listed in Table 4. We found that the SVM machine learning model provides best prediction values (based on MCC and accuracy values) for the RFE-*t*-test-SVM feature set.

**Table 4.**
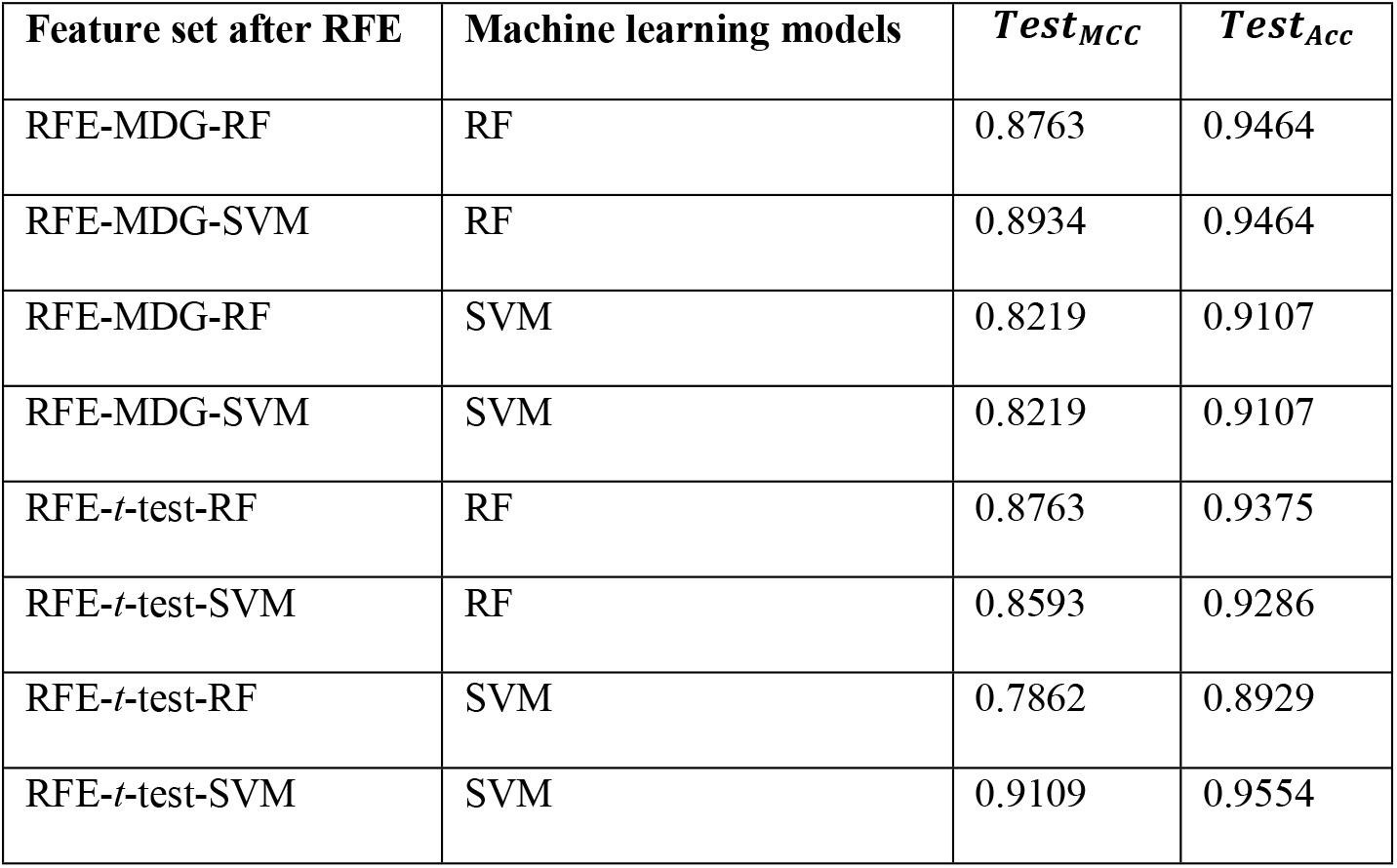
MCC and accuracy values obtained from RF and SVM for testing data for different RFE feature subsets.

| Feature set after RFE | Machine learning models | $Test_{MCC}$ | $Test_{Acc}$ |
| --- | --- | --- | --- |
| RFE-MDG-RF | RF | 0.8763 | 0.9464 |
| RFE-MDG-SVM | RF | 0.8934 | 0.9464 |
| RFE-MDG-RF | SVM | 0.8219 | 0.9107 |
| RFE-MDG-SVM | SVM | 0.8219 | 0.9107 |
| RFE-*t*-test-RF | RF | 0.8763 | 0.9375 |
| RFE-*t*-test-SVM | RF | 0.8593 | 0.9286 |
| RFE-*t*-test-RF | SVM | 0.7862 | 0.8929 |
| RFE-*t*-test-SVM | SVM | 0.9109 | 0.9554 |

### Performance comparison

Our best machine learning model’s prediction performance was compared to the sequence matching strategy-BLASTP [8, 9]. To identify bacteriocins sequences, BLASTP takes positive sequences of the training set as subject sequences and positive sequences of the testing set as query sequences and estimates the sequence similarity (percent identity) for each query sequence by aligning them with the subject sequences. Similarly, to detect non-bacteriocin sequences from BLASTP, we considered all negative sequences of the training and testing sets as subject and query sequences, respectively. Fig 5 shows the number of true positives and negatives with respective percent identity threshold for BLASTP tool. According to **S8 Table**, our best classifier SVM model with RFE-*t*-test-SVM has 53 true positives and 54 true negatives. BLASTP can identify a similar number of true positives and true negatives as our best classifier if we set the percent identify threshold of BLASTP lower than 30 and 20 for finding the true positives and true negatives, respectively. However, setting such a low percent identify threshold in BLASTP is very unrealistic and will increase false positive and false negative results.

**Fig 5.**
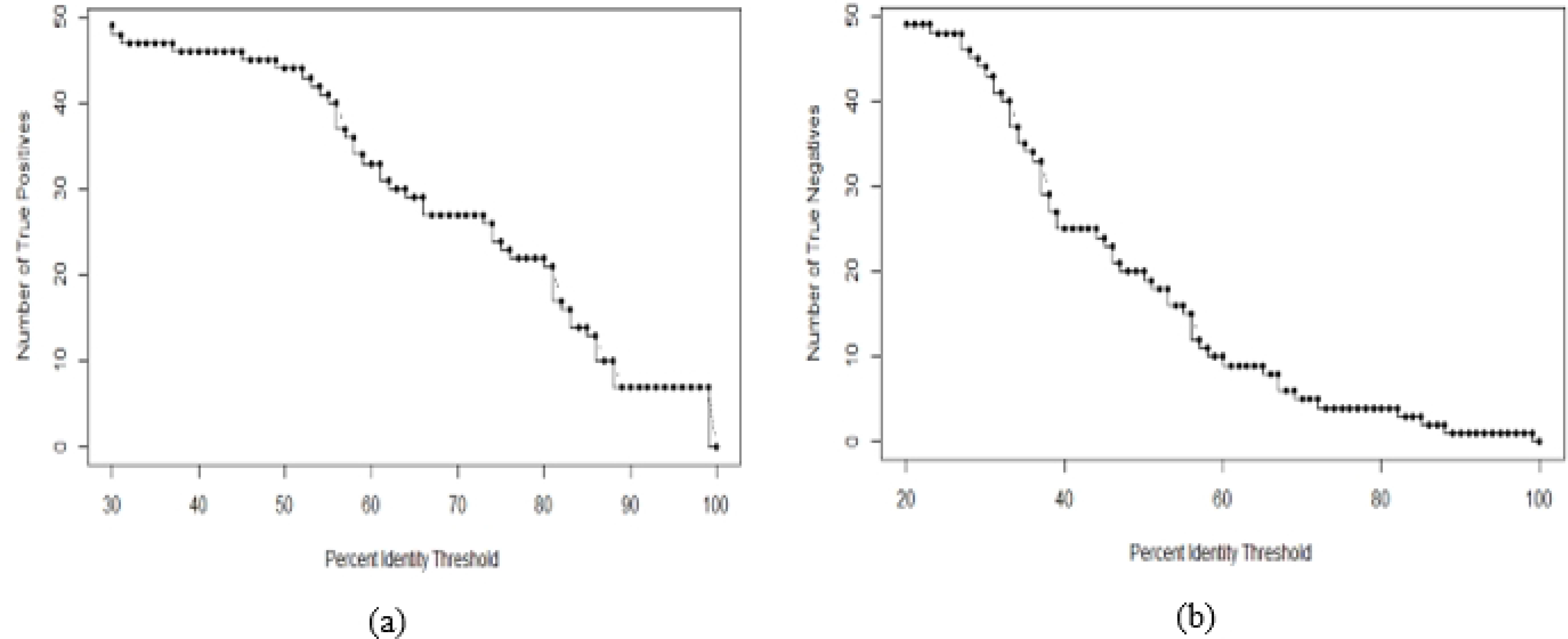
Identification of test sequences using BLASTP as a function of percent identity threshold (a) using bacteriocin sequences from the training data and (b) using non-bacteriocin sequences from the training data.

We also compared the performance of our method with a recent deep learning-based approach RMSCNN [13] developed for the bacteriocin prediction. RMSCNN takes positive and negative training protein sequences in FASTA format as inputs, encodes all amino acids of each protein sequences to some numbers defined in a protein dictionary, then constructs a matrix of the encoded sequences. This matrix is used to train a convolutional neural network where a random model is used to modify the scale of the convolutional kernel. To compare the prediction accuracy and runtime with our BacPred software tool, we ran RMSCNN with the same training and testing datasets that we used in our machine learning models. The runtime of RMSCNN or BacPred is defined as the total time required in training and testing phases. Note that our BacPred software was implemented based on our best model discussed earlier, that is, the SVM machine learning model with RFE-*t*-test-SVM feature set. Both RMSCNN and BacPred were executed in a machine with macOS operating system, 2.3 GHz 8-Core Intel Core i9 processor, and 32 GB 2667 MHz DDR4 memory configuration. Table 5 shows the prediction accuracy and runtime of both methods/tools, and our BacPred outperforms RMSCNN in terms of both accuracy and runtime.

**Table 5.** Accuracy and runtime (in seconds) of RMSCNN and BacPred.

| Method/tool | $Test_{Acc}$ | Runtime (sec.) |
| --- | --- | --- |
| RMSCNN | 0.9375 | 2007.86 |
| BacPred | 0.9554 | 217.84 |

Based on Table 5, we can infer that our method was able to identify the most important features to detect highly diverse bacteriocin sequences with higher accuracy and lower runtime. Researchers can easily use our optimal feature-based software tool to discover novel bacteriocin sequences. Since our software tool is open source, they can modify our tool to fit it in similar or completely new biological applications.

## Conclusion

Discovery of new bacteriocins is crucial to develop new antibiotic drugs to combat against antibiotic resistance. In this paper, we presented a machine learning-based approach for identifying novel bacteriocins. We extracted the applicant features from the primary and secondary attributes of protein sequences and then we analyzed all features based on Pearson correlation coefficient, *t*-test, and MDG values. We obtained two reduced feature sets of 140 and 44 features, and we further filtered out features using RFE technique. The final selected feature sets were considered as optimal sets of features and used to build the SVM and RF machine learning models. We found that SVM shows better prediction performance with the RFE-*t*-test-SVM-reduced feature set. The performance of our best model is compared to that of the sequence matching-based tool BLASTP. For BLASTP to obtain true positive as well as true negative results comparable to our best model requires a percent identity threshold so low that it is impractical. Also, our method showed better prediction accuracy with lower runtime compared to a deep learning-based method RMSCNN.

We also implemented a software package BacPred based on our best model to identify bacteriocin sequences by integrating all necessary tools and programs required for generating the optimal set of features automatically. Using our software tool, users will be able to predict unseen testing data for bacteriocin detection and can include new known bacteriocin and non-bacteriocin sequences to train data that eventually improve the predictive power of the machine learning model. Currently, our model is suitable to identify single bacteriocin protein sequence and we plan to update it to discover protein clusters of tailocins i.e., phage tail-like bacteriocins [28, 29]. Also, in the future, we will consider better characterized features such as position specific scoring matrix [30] and develop a more robust feature selection algorithm for better characterized features to increase the prediction accuracy of the machine learning models. Whenever more bacteriocin sequences are available, we will retain our model.

## Supporting information

**S1 File.** Training dataset composed of known bacteriocin and non-bacteriocin protein sequences.

(FASTA)

**S2 File.** Testing dataset composed of known bacteriocin and non-bacteriocin protein sequences.

(FASTA)

**S1-S8 Tables.** Confusion matrixes of the machine learning models.

(DOCX)

## Notes

### Competing Interest Statement

The authors have declared no competing interest.

## References

1. Control CfD, Prevention. Antibiotic resistance threats in the United States, 2019: US Department of Health and Human Services, Centres for Disease Control and Prevention; 2019.

2. Fields FR, Lee SW, McConnell MJ. Using bacterial genomes and essential genes for the development of new antibiotics. Biochemical pharmacology. 2017;134:74–86.

3. Riley MA, Wertz JE. Bacteriocins: evolution, ecology, and application. Annual Reviews in Microbiology. 2002;56(1):117–37.

4. Fields FR, Freed SD, Carothers KE, Hamid MN, Hammers DE, Ross JN, et al. Novel antimicrobial peptide discovery using machine learning and biophysical selection of minimal bacteriocin domains. Drug Development Research. 2020;81(1):43–51.

5. Hamid MN, Friedberg I. Bacteriocin detection with distributed biological sequence representation.

6. Van Heel AJ, de Jong A, Montalban-Lopez M, Kok J, Kuipers OP. BAGEL3: automated identification of genes encoding bacteriocins and (non-) bactericidal posttranslationally modified peptides. Nucleic acids research. 2013;41(W1):W448–W53.

7. Hammami R, Zouhir A, Le Lay C, Ben Hamida J, Fliss I. BACTIBASE second release: a database and tool platform for bacteriocin characterization. Bmc Microbiology. 2010;10(1):1–5.

8. Johnson M, Zaretskaya I, Raytselis Y, Merezhuk Y, McGinnis S, Madden TL. NCBI BLAST: a better web interface. Nucleic acids research. 2008;36(suppl_2):W5–W9.

9. Boratyn GM, Camacho C, Cooper PS, Coulouris G, Fong A, Ma N, et al. BLAST: a more efficient report with usability improvements. Nucleic acids research. 2013;41(W1):W29–W33.

10. Morton JT, Freed SD, Lee SW, Friedberg I. A large scale prediction of bacteriocin gene blocks suggests a wide functional spectrum for bacteriocins. BMC bioinformatics. 2015;16(1):1–9.

11. Hamid M-N, Friedberg I. Identifying antimicrobial peptides using word embedding with deep recurrent neural networks. Bioinformatics. 2019;35(12):2009–16.

12. Mikolov T, Chen K, Corrado G, Dean J. Efficient estimation of word representations in vector space. arXiv preprint arXiv:13013781. 2013.

13. Cui Z, Chen Z-H, Zhang Q, Gribova VV, Filaretov VF, Huang D-s. RMSCNN: A Random Multi-Scale Convolutional Neural Network for Marine Microbial Bacteriocins Identification. IEEE/ACM Transactions on Computational Biology and Bioinformatics. 2021.

14. O’Shea K, Nash R. An introduction to convolutional neural networks. arXiv preprint arXiv:151108458. 2015.

15. Gu J, Wang Z, Kuen J, Ma L, Shahroudy A, Shuai B, et al. Recent advances in convolutional neural networks. Pattern recognition. 2018;77:354–77.

16. Kononenko I, editor Estimating attributes: Analysis and extensions of RELIEF. European conference on machine learning; 1994: Springer.

17. Robnik-Šikonja M, Kononenko I, editors. An adaptation of Relief for attribute estimation in regression. Machine learning: Proceedings of the fourteenth international conference (ICML’97); 1997.

18. Leo B. Random forests. Machine learning. 2001;45(1):5–32.

19. Fu L, Niu B, Zhu Z, Wu S, Li W. CD-HIT: accelerated for clustering the next-generation sequencing data. Bioinformatics. 2012;28(23):3150–2.

20. Bhasin M, Raghava GP. Classification of nuclear receptors based on amino acid composition and dipeptide composition. Journal of Biological Chemistry. 2004;279(22):23262–6.

21. Chou KC. Prediction of protein cellular attributes using pseudo‐amino acid composition. Proteins: Structure, Function, and Bioinformatics. 2001;43(3):246–55.

22. Chou K-C. Using amphiphilic pseudo amino acid composition to predict enzyme subfamily classes. Bioinformatics. 2005;21(1):10–9.

23. Dubchak I, Muchnik I, Holbrook SR, Kim S-H. Prediction of protein folding class using global description of amino acid sequence. Proceedings of the National Academy of Sciences. 1995;92(19):8700–4.

24. Calle ML, Urrea V. Stability of Random Forest importance measures. Briefings in bioinformatics. 2011;12(1):86–9.

25. Chowdhury AS, Reehl SM, Kehn-Hall K, Bishop B, Webb-Robertson B-JM. Better understanding and prediction of antiviral peptides through primary and secondary structure feature importance. Scientific reports. 2020;10(1):1–8.

26. Chicco D, Jurman G. The advantages of the Matthews correlation coefficient (MCC) over F1 score and accuracy in binary classification evaluation. BMC genomics. 2020;21(1):1–13.

27. Chicco D, Tötsch N, Jurman G. The Matthews correlation coefficient (MCC) is more reliable than balanced accuracy, bookmaker informedness, and markedness in two-class confusion matrix evaluation. BioData mining. 2021;14(1):1–22.

28. Patz S, Becker Y, Richert-Pöggeler KR, Berger B, Ruppel S, Huson DH, et al. Phage tail-like particles are versatile bacterial nanomachines–A mini-review. Journal of advanced research. 2019;19:75–84.

29. Ghequire MG, De Mot R. The tailocin tale: peeling off phage tails. Trends in microbiology. 2015;23(10):587–90.

30. Guigo R. An introduction to position specific scoring matrices. Bioinformatica upf edu. 2016.

